# Canagliflozin increases adenoma burden in female APC^Min/+^ mice

**DOI:** 10.1101/2021.03.27.437278

**Authors:** Justin Korfhage, Mary E. Skinner, Jookta Basu, Joel K. Greenson, Richard A. Miller, David B. Lombard

**Author notes:** Correspondence: David B. Lombard, M.D., Ph.D., University of Michigan, 3015 BSRB, 109 Zina Pitcher Place, Ann Arbor, MI 48109.

## Abstract

The diabetes drug canagliflozin acts primarily by inhibiting glucose reuptake by the sodium glucose transporter 2 (SGLT2) in the kidney proximal tubule, thereby lowering serum glucose levels. Canagliflozin also acts on SGLT1, a related transporter responsible for glucose uptake in the small intestine and more distal kidney tubules. Several cancers overexpress SGLT1 and SGLT2, where these transporters fuel tumor metabolism. A recent study by NIA’s Interventions Testing Program (ITP) showed that canagliflozin treatment extends lifespan in male mice. Since cancer is the major cause of death in most mouse strains, including the UM-HET3 strain used by the ITP, this observation suggests that canagliflozin might exert anti-cancer effects in this context. Here, we treated a commonly-used mouse neoplasia model -- the intestinal adenoma-prone APC^Min/+^ strain -- with canagliflozin, to test the effects of drug treatment on tumor burden. Surprisingly, canagliflozin increased the total area of intestine involved by adenomas, an effect that was most marked in the distal intestine and in female mice. Immunohistochemical analysis suggested that canagliflozin may not influence adenoma growth via direct SGLT1/2 inhibition in neoplastic cells themselves. Instead, our results are most consistent with a model whereby canagliflozin aggravates adenoma development by altering the anatomic distribution of intestinal glucose absorption, as evidenced by increases in postprandial GLP-1 levels consistent with delayed glucose absorption. Our results suggest that canagliflozin exacerbates adenomatosis in the APC^Min/+^ model via complex, cell-non-autonomous mechanisms, and hint that sex differences in incretin responses may underlie differential effects of this drug on lifespan.

## Introduction

Canagliflozin is a member of the gliflozin class of SGLT2 inhibitors used to treat type 2 diabetes mellitus (Nomura et al., 2010). SGLT2 inhibitors bind the sodium glucose symporter to inhibit renal glucose reuptake in the proximal tubule, leading to increased urinary glucose excretion and reduced blood glucose levels (Tsujihara et al., 1999). To a lesser degree, canagliflozin also acts on SGLT1, a related sodium glucose symporter. SGLT1 and SGLT2 are partially functionally redundant in the kidney (Grempler et al., 2012). SGLT1 is also expressed on the apical membrane of the small intestinal epithelium, where it functions as the primary route of glucose absorption in the gastrointestinal (GI) tract (Gorboulev et al., 2012; Sala-Rabanal et al., 2018). Canagliflozin is unique among FDA-approved SGLT2 inhibitors, in that it has a relatively low selectivity (roughly 200-fold) for SGLT2 versus SGLT1, as opposed to other members of the gliflozin class of drugs which are far more selective for SGLT2 (Ohgaki et al., 2016). Canagliflozin, administered orally, is present at much higher concentrations in the GI tract than in plasma (Oguma et al., 2015). Plasma concentrations of canagliflozin elicited by clinical dosing regimens (approximately 5 μM) are insufficient to inhibit SGLT1 (Sha et al., 2015). However, the high small intestine luminal canagliflozin concentrations observed after oral canagliflozin administration are sufficient to inhibit intestinal SGLT1-mediated glucose uptake in rodents and humans (Kuriyama et al., 2014; Oguma et al., 2015; Ohgaki et al., 2016; Polidori et al., 2013), leading to increased luminal glucose levels in the distal small intestine and colon. Such changes in luminal contents can influence release of GI hormones called incretins. Several studies have demonstrated that SGLT1 inhibition or knockout delays intestinal glucose absorption, thereby stimulating the second phase of biphasic release of the incretin GLP-1, a hormone that regulates satiety, gastric emptying, intestinal growth, and glucose-mediated insulin secretion (Kuriyama et al., 2014; Oguma et al., 2015; Paternoster & Falasca, 2018; Polidori et al., 2013).

There has been great interest in effects of SGLT2 inhibitors on neoplasia and in repurposing these drugs to treat cancer and other diseases. An initial study of the carcinogenicity of canagliflozin in rats found an increase in the incidence of pheochromocytoma as, renal tubular tumors, and Leydig cell tumors in canagliflozin-treated animals; however, somewhat paradoxically, overall lifespan was increased in male (and to a much lesser extent, female) rats administered canagliflozin (De Jonghe et al., 2014). Follow-up studies attributed the renal tumors to carbohydrate malabsorption (De Jonghe et al., 2017; Mamidi et al., 2014). Likewise, a recent study by the National Institute of Aging’s (NIA) Interventions Testing Program (ITP) demonstrated an increased lifespan in male mice treated with canagliflozin (Miller et al., 2020). Since most mice in the UM-HET3 strain background used by the ITP die of cancer (Lipman, Galecki, Burke, & Miller, 2004), these results suggest that canagliflozin may inhibit cancer development and/or progression in male mice of this strain. In this regard, a recent meta-analysis revealed that adults with type 2 diabetes who take canagliflozin exhibited a significantly lower incidence of GI cancers relative to controls (Tang et al., 2017). Several studies have demonstrated overexpression of SGLT1 or SGLT2 in many tumor types, where these transporters import glucose for energy production and provide carbon skeletons for macromolecular biosynthesis (Koepsell, 2017). SGLT1 is overexpressed in colorectal carcinoma, and that its expression levels correlate with disease stage (Guo et al., 2011). Canagliflozin and other SGLT2 inhibitors decrease cancer cell proliferation in cell culture and tumor growth in xenograft and genetically engineered mouse cancer models (Hung et al., 2019; Kaji et al., 2018; Kawaguchi et al., 2019; Scafoglio et al., 2015; Shiba et al., 2018; Villani et al., 2016). Overall, these studies suggest that canagliflozin may exert complex, species- and tumor type-specific effects on tumorigenesis.

The APC^Min/+^ mouse is a commonly used model of GI adenomatosis. This strain carries a heterozygous germline mutation in the adenomatous polyposis coli (*APC*) gene (Moser, Pitot, & Dove, 1990), mimicking the human condition Familial Adenomatous Polyposis (FAP). The protein product of the wildtype *APC* gene suppresses proliferative signaling via the WNT-beta Catenin axis (Klaus & Birchmeier, 2008). Upon loss of the remaining wildtype APC allele, an intestinal adenoma forms; in humans, these lesions possess the potential for additional genetic changes and malignant progression. Humans carrying *APC* mutations develop numerous colonic adenomas and have a high predisposition for developing colorectal carcinoma (Kinzler & Vogelstein, 1996). APC^Min/+^ mice develop primarily small intestinal adenomas, leading to lethality around 120 days of age, typically due to intestinal obstruction and bleeding (Moser et al., 1990; Su et al., 1992). Importantly, studies have shown that two other drugs with longevity-extending effects in the UM-HET3 strain, acarbose and rapamycin, likewise attenuate adenomatosis in APC^Min/+^ mice (Dodds et al., 2020; Hasty et al., 2014; Quesada et al., 1998). Given the protective effects of canagliflozin against human GI cancers and the lifespan extension observed with canagliflozin treatment in rodents, here we evaluated the effects of canagliflozin treatment on adenoma development in APC^Min/+^ mice.

## Results

### Canagliflozin treatment increases adenoma burden in APC^Min/+^ mice

To test potential effects of canagliflozin on adenoma development in APC^Min/+^ mice, animals were fed canagliflozin-containing chow from 36 days of age until the point of sacrifice at 106-108 days of age, using a canagliflozin concentration previously shown to extend lifespan in male UM-HET3 mice (Miller et al., 2020). Prior to sacrifice, mice were fasted for 18 hours to minimize local variability in protein expression due to intestinal contents (Figure 1A). We observed a sex specific effect: males, but not females, treated with canagliflozin were significantly lighter than controls (p_sex:cana_ = 0.048; Figure S1A). No significant differences were observed in fasted blood glucose levels in male or female mice treated with canagliflozin (Figure S1B).

**Figure 1.**
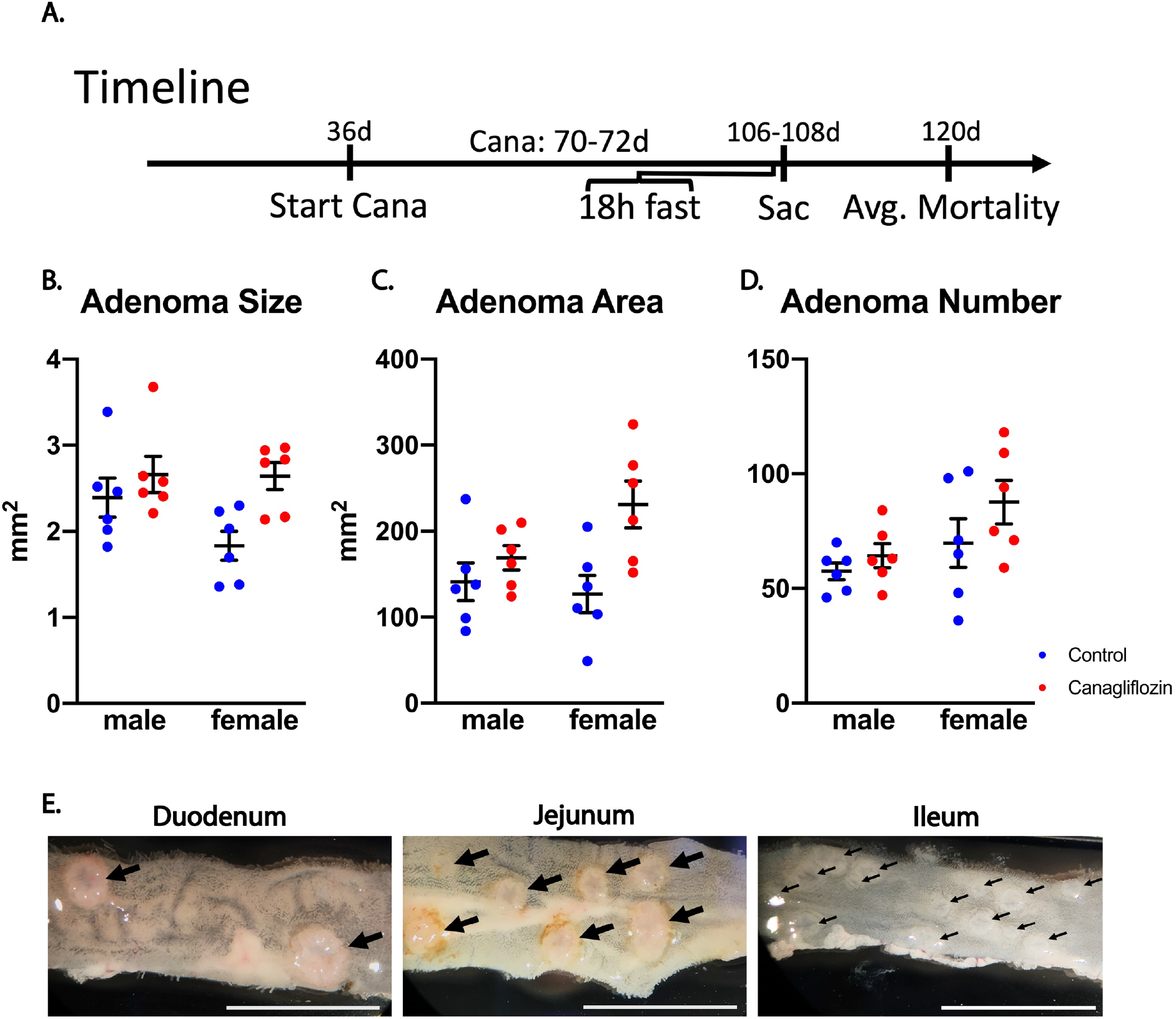
Canagliflozin Increases Adenoma Burden in APC^Min/+^ Mice. **A)** Timeline of canagliflozin treatment and feeding procedures of APC^Min/+^ mice prior to sacrifice (sac). Mice were fed a diet of canagliflozin (cana) or control chow for 70-72 days then fasted for 18 hours prior to sacrifice. **B)** Average cross-sectional area of adenomas from control and canagliflozin treated mice was measured using ImageJ oval tool. Each point represents the mean adenoma size from a single mouse. Adenomas in canagliflozin treated mice (mean: 2.631 mm^2^) were significantly larger than adenomas in control mice (mean: 2.114 mm^2^); drug effect, p < 0.05, two-way ANOVA; sex effect and interaction were n.s. **C)** The total area of the small intestine involved by adenomas in each mouse was summed and plotted. Each point represents an individual mouse. The area of the small intestine involved by adenomas in canagliflozin mice (mean: 200 mm^2^) was significantly greater than control mice (mean: 134 mm^2^); drug effect, p < 0.01, two-way ANOVA; sex effect and interaction were n.s. **D)** The number of adenomas in each mouse was counted and plotted. Each point represents one mouse. Canagliflozin-treated mice (mean: 76) did not exhibit significantly more adenomas than control mice (mean: 63.67); drug effect, p = 0.22, two-way ANOVA; sex effect and interaction were n.s. **E)** Representative images of adenomas in the duodenum, jejunum, and ileum. Adenomas are identified with black arrows. Scalebar = 10 mm.

To determine the effects of canagliflozin on adenoma development, the small intestine was dissected into thirds, approximating the duodenum, jejunum, and ileum. The entire length of each section was imaged, and adenomas were counted and measured. A 1.3-fold increase in adenoma size was observed in canagliflozin-treated mice compared to controls (p_cana_ = 0.011; Figure 1B). The overall adenoma area was then calculated for each mouse. Canagliflozin-treated mice had a 1.5-fold greater total area of intestine involved by adenomas (p_cana_ = 0.007; Figure 1C). There was a modest, non-significant increase in the number of adenomas present per mouse (p_cana_ = 0.22; Figure 1D).

We observed marked differences in adenoma size and number across different regions of the small intestine, consistent with previous studies in which adenomas forming in the APC^Min/+^ background were observed to be smaller, but more numerous, in the distal small intestine (Figure 1E) (Koehler et al., 2015; Moser et al., 1990). By histologic analysis, there was no evidence of tissue invasion or progression to carcinoma in any adenomas examined (Figure S1C). Taken together, these results suggest that 1) increases in adenoma burden associated with canagliflozin treatment are mostly due to augmentation of adenoma growth rather than increases in adenoma number, and 2) canagliflozin exerts no obvious effects on the grade or stage of lesions in the APC^Min/+^ mouse.

### Effects of canagliflozin on intestinal adenomas exhibit evidence of sex and region specificity

Based on the male-specific lifespan extension observed with this canagliflozin dosage in UM-HET3 mice (Miller et al., 2020) and the fact that the three intestinal regions are histologically and physiologically distinct, we analyzed adenoma number, adenoma area, and average adenoma size to assess the effects of canagliflozin, sex, intestinal region, or interactions on these metrics. There was a significant interaction between canagliflozin and region in adenoma size, suggesting that canagliflozin exerted its effects on adenoma size most prominently in the distal small intestine (p_cana:region_ = 0.0002; Figure 2A), and a nearly significant interaction between these two variables in total adenoma area (p_cana:region_ = 0.054; Figure 2B). Overall, the magnitude of the effects of canagliflozin on adenoma size and area were greater toward the distal small intestine in females; however, these three-way interactions were not statistically significant (Figure 2 A-B, S2 A-B). Adenoma size and total adenoma area were not significantly influenced by canagliflozin in a sex specific manner (Figure 2 A-B), however there were significant increases in adenoma size and total adenoma area associated with canagliflozin treatment in females (Student’s t-test; Figure 2 A-C). Males exhibited smaller and less numerous adenomas in each intestinal region than females by every metric (Figure 2 A-C). These data suggest that chronic canagliflozin treatment increases adenoma size and area in the distal small intestine, changes that are significant in female mice.

**Figure 2.**
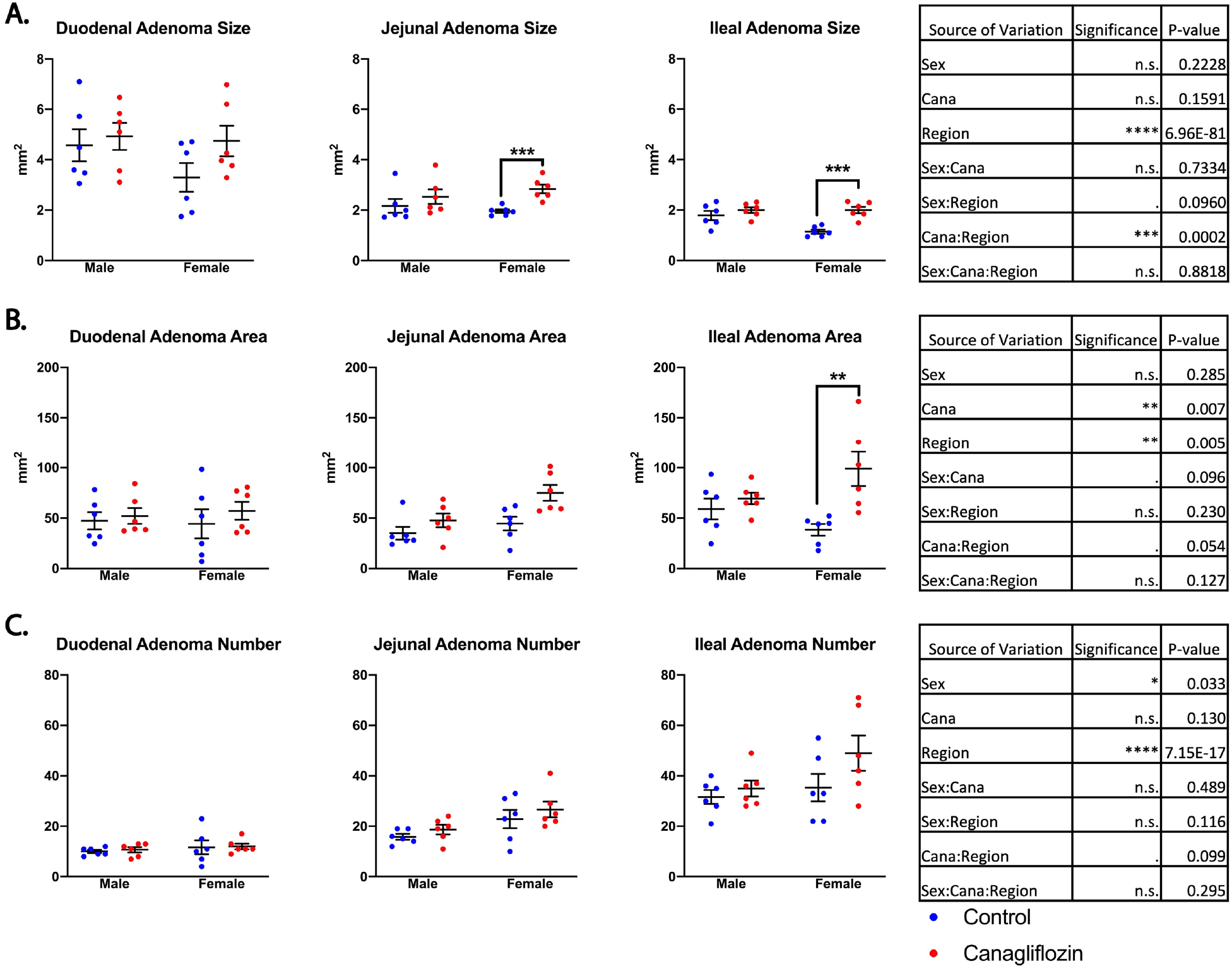
Increased Adenoma Burden in Canagliflozin Treated Mice is Concentrated in the Distal Small Intestine. **A)**The variance in the size of individual adenomas accounted for by canagliflozin (cana), sex, and region was analyzed by a linear mixed effects model, treating each mouse ID as a random effect and region as a repeated measure. Mean adenoma size was plotted for each mouse. **B)** Total cross-sectional area involved by adenomas was plotted for each mouse. 3-way ANOVA was performed using a linear mixed effects model with region as a repeated measurement. **C)** Adenoma number in each region of the small intestine was analyzed for each mouse and compared by 3-way ANOVA using a linear mixed effects model with region as a repeated measurement. **A-C)** Each point represents one mouse. ANOVA tables are displayed to the right of each figure. Significant differences within sex and region were determined by two tailed Student’s t test and are displayed on each graph. * p < 0.05, ** p < 0.01, *** p < 0.001, **** p < 0.0001.

### Proteins involved in SGLT glucose import are mis-localized in adenomas

Significant canagliflozin-induced increases in tumor burden in female mice were unanticipated, particularly since this drug increases lifespan in male mice and rats *(De Jonghe et al*., *2014; Miller et al*., *2020)*. We hypothesized that these differences in response to canagliflozin treatment might be related to sex differences in expression of glucose transporters in the ileum. In this regard, Oguma *et al*. reported that transient SGLT1 inhibition with orally administered canagliflozin decreased duodenal and jejunal carbohydrate absorption and increased luminal carbohydrate content in rats (Oguma et al., 2015). To address this issue, we stained intestinal tissue sections for proteins involved in intestinal glucose import. The luminal side of the intestinal epithelium expresses SGLT1, which exploits the sodium concentration gradient to promote import of glucose against its concentration gradient. Sodium export via the Na^+^/K^+^ ATPase at the basolateral membrane of the intestinal epithelium maintains a sodium concentration gradient to allow SGLT1 to import sodium and glucose. GLUT2 serves as a channel for glucose efflux on the basolateral membrane of the intestinal epithelium (Thorens & Mueckler, 2009).

We performed immunohistochemical staining for these proteins in the ilea of male and female mice to test for potential sex differences in expression and expression changes with canagliflozin treatment. We also stained for SGLT2 to test the possibility that intestinal adenomas may aberrantly express SGLT2. No expression of SGLT2 on the luminal membrane of the intestine was observed (Figure 3A). Likewise, SGLT2 expression was not detected on adenomas, suggesting that any effects of canagliflozin in this model are unlikely mediated via intestinal SGLT2 (Figure 3A). As a positive staining control, strong SGLT2 staining on the apical membrane of the kidney proximal tubule was readily detectable under the same staining conditions (Figure S3). As expected, SGLT1 staining was ubiquitous across the entire apical surface of the ileal epithelium (Figure 3A). Adenomas exhibited a significant decrease in apical membrane SGLT1 expression relative to the ileal epithelium, with almost no detectable membrane expression within the adenoma (p_adenoma_ = 0.017; Figure 3A-B). The low membrane SGLT1 expression in adenomas renders it unlikely that the effects of canagliflozin on adenoma growth represent a direct effect of the drug on SGLT1 in the tumors themselves, although we cannot rule out the possibility that SGLT1 is expressed at a level below the sensitivity of immunohistochemical staining. Furthermore, expression of the Na^+^/K^+^ ATPase was significantly decreased on the membranes of adenomas relative to the surrounding tissue (p_adenoma_ < 0.0001; Figure 3C). Adenomas exhibited significantly higher levels of GLUT2 staining relative to the surrounding tissue, which could allow for passive glucose transport to fuel the growth of neoplastic tissue (p_adenoma_ = 0.076; Figure 3D). These data suggest that the pro-neoplastic effects of canagliflozin in the APC^Min/+^ model are unlikely to reflect direct action on neoplastic cells, and that very low expression of SGLT1/2 may be compensated for by increases in GLUT2 expression within the adenoma.

**Figure 3.**
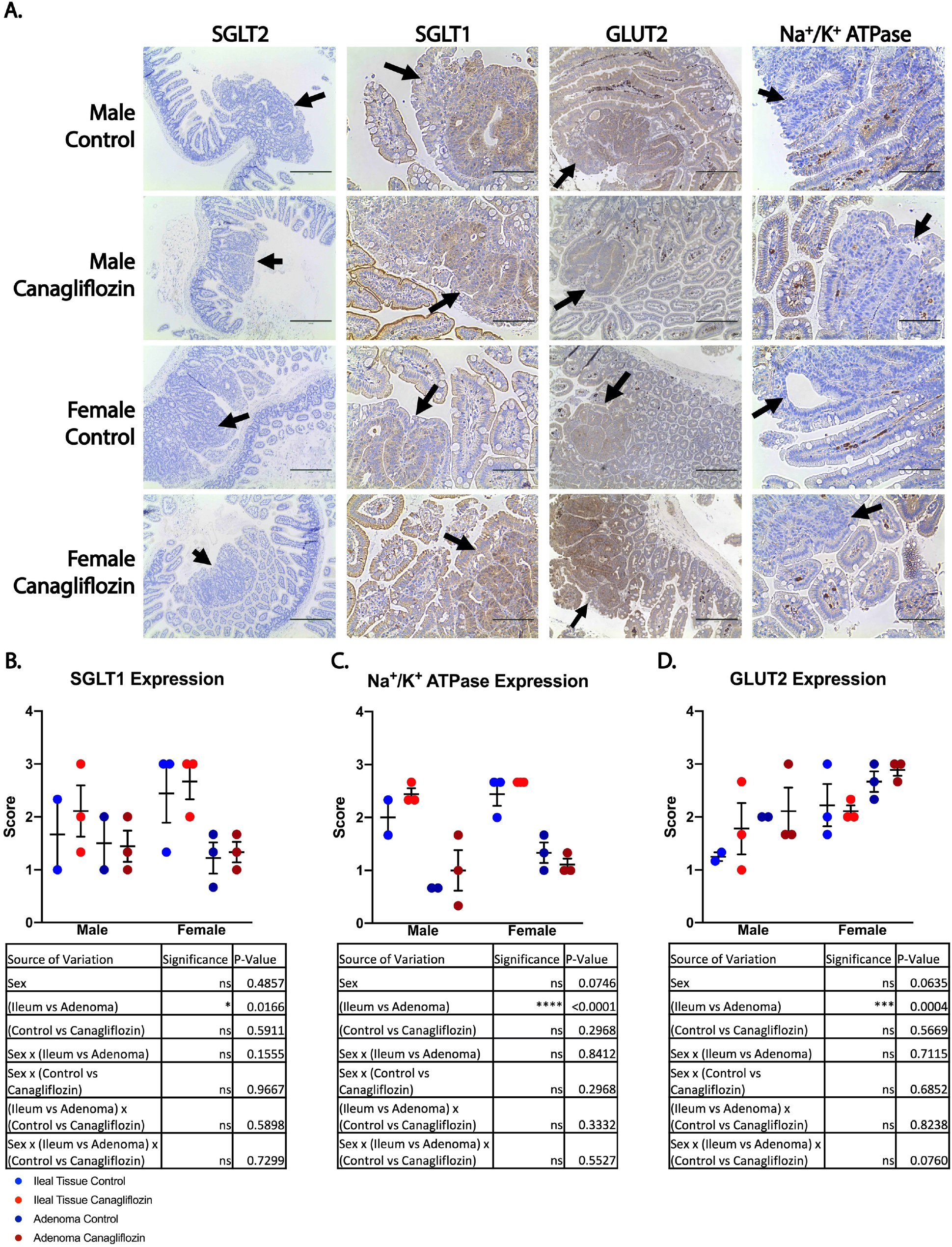
Adenomas Exhibit Decreased Membrane Expression of Canonical Intestinal Glucose Import Machinery. **A)**Representative images of immunohistochemical staining for proteins relevant to intestinal glucose transport. Adenomas are indicated with arrows; SGLT2 scalebar = 400 μm, SGLT1 scalebar = 100 μm, GLUT2 scalebar = 200 μm, Na^+^/K^+^ ATPase scalebar = 100 μm. **B-D)** expression scoring from membranes of neoplastic cells and normal tissue. Each point represents the mean score from 3 images from the ileum of one mouse. The contributions of sex, canagliflozin, and normal ileal tissue vs neoplastic tissue were analyzed by 3-way ANOVA. ANOVA tables are presented below the figure. * p < 0.05, *** p < 0.001, **** p < 0.0001.

### Canagliflozin elevates postprandial GLP-1 levels

Pharmacologic inhibition or genetic knockout of SGLT1 delays postprandial glucose absorption and increases postprandial plasma GLP-1 levels (Gorboulev et al., 2012; Oguma et al., 2015; Powell et al., 2013; Shibazaki et al., 2012; Zambrowicz et al., 2012). Circulating active GLP-1 suppresses appetite, augments insulin release, and slows gastric emptying, among other effects (Paternoster & Falasca, 2018). Active GLP-1 is rapidly cleaved to its inactive form by DPP4. GLP-1 also exerts intestinotrophic effects; endogenous GLP-1 induces intestinal growth, and GLP-1 receptor activation stimulates adenoma formation in APC^Min/+^ mice (Kissow et al., 2012; Koehler et al., 2015; Simonsen et al., 2007). We therefore measured serum GLP-1 and active GLP-1 levels in a separate set of canagliflozin-treated and control mice as an indirect measure of intestinal SGLT1 inhibition, to determine whether increased GLP-1 induced by canagliflozin treatment might represent a potential driver of adenoma growth.

Based on prior studies of mouse GI tract transit time, serum samples were harvested three hours after fasting and refeeding in order to obtain serum at the time when the chow was transiting the small intestine (Padmanabhan, Grosse, Asad, Radda, & Golay, 2013). This experiment revealed a significant canagliflozin-dependent increase in total GLP-1 (p_cana_ = 0.007; Figure 4A). There was a suggestion that the canagliflozin effect was specific to males, but this did not reach statistical significance (p_cana_ = 0.068; Figure 4A). However, there was a significant increase in total GLP-1 with canagliflozin treatment in males (Student’s t-test; Figure 4B). Females also exhibited significantly higher levels of total GLP-1 than males (p_sex_ = 0.038; Figure 4A). Levels of active GLP-1 showed a similar pattern, in which males displayed a greater canagliflozin-dependent increase, though none of these effects were statistically significant (Figure 4C). Unlike the levels of total GLP-1, active GLP-1 levels were not higher in control females than control males (Figure 4C). Across the entire population, there was a trend for canagliflozin treatment to increase active GLP-1 levels (p_cana_ = 0.07; Figure 4C). With the caveat that the sample size examined here was relatively modest, these data support the hypothesis that canagliflozin administration shifts glucose absorption to the more distal small intestine, thus driving increased production of GLP-1.

**Figure 4.**
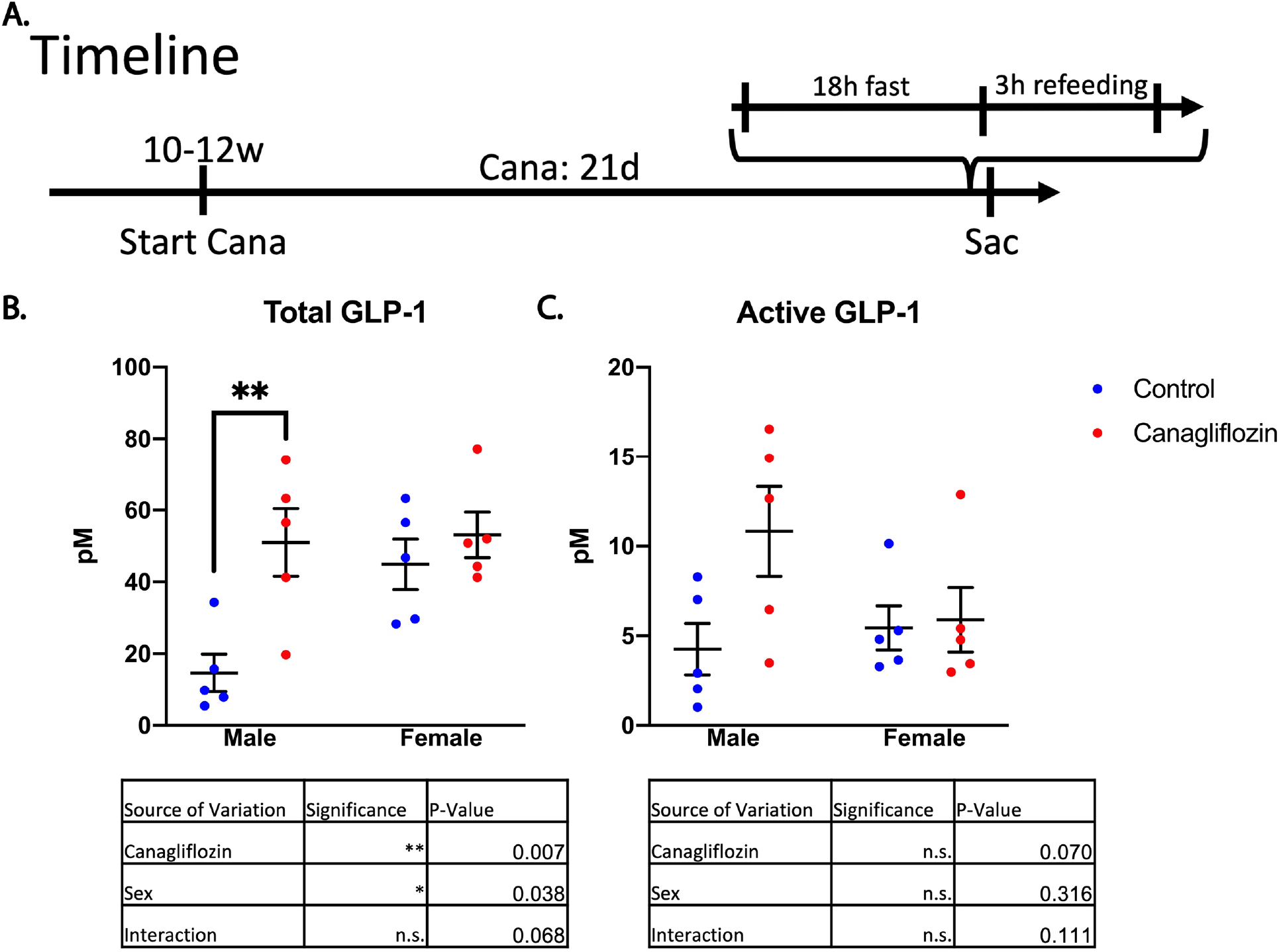
Canagliflozin Treatment Elevates Postprandial Serum GLP-1 levels. **A)**Timeline of canagliflozin treatment prior to obtaining serum. **B)** Total GLP-1 levels were measured by colorimetric ELISA assay. Each point represents the mean of two technical replicates from the serum of one mouse. **C)** Active GLP-1 levels were measured by a luminescent ELISA assay. Each point represents the mean of two technical replicates from the serum of one mouse. **B-C)** The effects of canagliflozin and sex were analyzed by 2-way ANOVA. ANOVA tables are presented below graphs. Significant differences within each sex were determined by two tailed Student’s t test and are displayed on each graph. * p < 0.05, ** p < 0.01, *** p < 0.001, **** p < 0.0001.

## Discussion

Canagliflozin, an SGLT2 inhibitor prescribed for the treatment of diabetes, suppresses tumor development in some animal model systems, decreases the incidence of GI cancer in diabetic patients, and extends lifespan in male mice and rats (De Jonghe et al., 2014; Hung et al., 2019; Kaji et al., 2018; Kawaguchi et al., 2019; Miller et al., 2020; Scafoglio et al., 2015; Shiba et al., 2018; Tang et al., 2017; Villani et al., 2016). Here, we tested the effects of canagliflozin administration on adenoma burden in the APC^Min/+^ mouse intestinal adenomatosis model, to begin to address the hypothesis that canagliflozin-mediated lifespan extension may be driven by delay or prevention of neoplasia. Surprisingly, canagliflozin-treated mice exhibited an increase in adenoma burden relative to controls. The overall increase in adenoma burden in our data was driven primarily by increases in adenoma size in the distal small intestine. There was evidence that these findings were female-specific, but the effect did not reach statistical significance in our sample size. Based on significant increases in adenoma size, but lack of significant differences in adenoma number between canagliflozin-treated mice and controls, it seems most likely that canagliflozin treatment primarily drives adenoma growth, rather than initiation. Crucially, our findings demonstrate that lifespan-extending interventions identified by the ITP do not necessarily have generalizable antineoplastic effects and that lifespan extension may occur in spite of tissue-specific, pro-neoplastic effects in specific cancer models. Superficially, these findings seem at odds with the protective effects of other lifespan-extending drugs in the APC^Min/+^ mouse model, specifically acarbose and rapamycin. Importantly, however, these studies either examined only males (Dodds et al., 2020), or did not analyze sexes separately (Hasty et al., 2014; Quesada et al., 1998).

Immunohistochemical analysis of cellular glucose import machinery expression in the intestine revealed no differences between males and females, or between canagliflozin-treated and control mice (Figure 3). Adenomas exhibited significantly reduced membrane expression of SGLT1 and the Na^+^/K^+^ ATPase relative to surrounding tissue, and they did not appear to localize to the apical and basolateral membranes respectively. Improper localization of these proteins crucial for intestinal glucose absorption could impair SGLT1-mediated glucose uptake. Conversely, GLUT2 was overexpressed in adenomas relative to the surrounding tissue suggesting that adenomas may employ this transporter to compensate for reduced SGLT1 expression to absorb sufficient quantities of glucose. This observation, along with lack of detectable SGLT2 expression in adenomas or normal intestine, suggested that increased adenomatosis in canagliflozin-treated mice may not be driven by direct effects of canagliflozin on the SGLT mediated glucose absorption in adenomas. Furthermore, direct effects of canagliflozin on neoplastic cells would be most likely to occur in the proximal small intestine, where luminal canagliflozin concentrations would be the highest prior to absorption by the small intestine. Instead, we observe the greatest increase in adenoma burden in the ileum, suggesting that this effect may be driven by downstream physiological effects of canagliflozin.

Increases in postprandial GLP-1 release are typical of SGLT1, but not SGLT2, inhibition or deficiency (Powell et al., 2013). Indeed, we observed increased postprandial GLP-1 levels in serum of canagliflozin-treated mice, similar to the effects of pharmacological inhibition or genetic knockout of SGLT1 (Gorboulev et al., 2012; Oguma et al., 2015; Powell et al., 2013; Shibazaki et al., 2012; Zambrowicz et al., 2012), suggesting that inhibition of intestinal SGLT1 was achieved using our canagliflozin administration regimen. Similar increases in GLP-1 are observed with acarbose treatment in human patients (McCarty & DiNicolantonio, 2015). This is thought to be caused by delay of glucose absorption and subsequent increase in delivery of glucose to the distal small intestine where it stimulates enteroendocrine L-cells to secrete GLP-1(Gribble & Reimann, 2019; McCarty & DiNicolantonio, 2015; Oguma et al., 2015). GLP-1 is a known intestinotrophic incretin (Kissow et al., 2012; Simonsen et al., 2007), and GLP-1 receptor activation drives adenoma growth in APC^Min/+^ mice (Koehler et al., 2015). However, we observe more prominent increases in adenoma burden in female mice, while total and active GLP-1 elevation was more prominent in male mice. Therefore, our findings may not support the hypothesis that aggravated adenoma development in this model is driven by GLP-1. Elevated GLP-1 levels may contribute to male-specific lifespan extension by canagliflozin and acarbose. GLP-1 ameliorates age associated pathologies in many tissue types; exerting cardioprotective effects, improving glucose tolerance, promoting weight loss, and protecting against neurodegeneration (McCarty & DiNicolantonio, 2015). Our observation of male specific weight loss induced by canagliflozin is consistent with the GLP-1 elevation we see in male canagliflozin treated mice (Figure S1A, 4B). We hypothesize that GLP-1 levels may be specifically elevated in acarbose-treated male mice, but not in females; to our knowledge, such measurements have not been performed.

Our findings underscore the need to understand the complex interactions between SGLT inhibitors, SGLT1, SGLT2, and GLP-1, and how these interactions may differ in males and females. SGLT2 inhibitors are already popular diabetes therapies; GLP-1 receptor agonists are FDA-approved, and dual SGLT1/2 inhibitors are being investigated as novel diabetes drugs. Our studies highlight the potential for unexpected sex-specific effects, and on-target side effects of such therapeutic approaches.

### Experimental procedures Canagliflozin Treatment

All vertebrate animal studies were performed in accord with animal protocol PRO00009721 (DL) and institutional IACUC policies. 24 C57BL/6J-Apc<Min>/J (002020) mice (12 males, 12 females) from Jackson Laboratories were fed *ad libitum* with chow containing 180 parts per million canagliflozin or with control chow starting at 36 days of age and continuing for 70 to 72 days. Food was replenished as needed. Prior to sacrifice, mice were fasted for 18 hours. At the end of the study, mice were euthanized over the course of 3 days. Each day, two mice from each group (male and female, +/-canagliflozin treatment) were randomly selected, humanely sacrificed, and dissected.

### Glucometer Reading

To obtain blood for glucose measurement, a shallow incision was made near the tail tip between 06:00 and 07:00 after an 18 h fast. A OneTouch Ultra blood glucose meter was used to obtain blood glucose measurements, taken in duplicate.

### Dissection and Tissue Preparation

Dissections were performed by an investigator blinded to drug treatment. Animals were anesthetized with isoflurane overdose, followed by organ removal. Kidneys were harvested and washed in PBS. One kidney from each mouse was preserved in 10% buffered formalin and the other was flash frozen in liquid nitrogen. Small intestines, extending from the pyloric sphincter to the ileocecal junction, were excised. Intestinal contents were flushed with 10-20 ml of ice-cold PBS. The small intestine was transferred to ice cold 10% buffered formalin for approximately 30 seconds. A single incision along the length of the small intestine was made to expose the luminal surface. The intestine was then divided into thirds, approximating the duodenum, jejunum, and ileum. These sections were imaged, rolled, and preserved in 10% buffered formalin.

### Adenoma Analysis

The full length of each intestinal section was imaged using a camera mounted to a dissecting microscope (Nikon SMZ645, C-W-10XB/22 eyepiece, 5x magnification). A ruler was included in each image to scale images in ImageJ to a 10 mm linear segment on the ruler. This scale was used to measure the approximate area of each adenoma using the ImageJ oval tool. Visible protrusions from the luminal face of the small intestine were counted as adenomas. All analyses were performed by an investigator blinded to the sex and drug treatment of the mice.

### Histology

Histologic staining was performed by the University of Michigan Department of Pathology Research Histology Core. The following antibodies were used in in histologic staining: SGLT2: Abcam anti-SGLT2 antibody (ab37296), 1:100 dilution; SGLT1: Abcam anti-SGLT1 antibody (ab14686), 1:100 dilution; Na^+^/K^+^ ATPase: Santa Cruz Biotechnology Na^+^/K^+^ ATPase α1 (C464.6): sc-21712, 1:1500 dilution.

### Scoring Protein Expression

Protein expression was scored on a scale from 0-3 by an individual blinded to sample identity, comparing staining intensity to mouse kidney tissue, which served as a positive control. Cytoplasmic, nuclear, and membrane staining was scored based on three representative images per mouse from ileal tissue sections containing regions of normal epithelium and neoplastic tissue. Images were captured blinded to the sample identity. The average of the three scores in each mouse was plotted. Analyses were performed by an investigator blinded to sample identities.

### GLP-1 Measurement

Ten male and ten female wild-type 10-12 week old C57BL/6J mice (000664) from Jackson Laboratories were fed canagliflozin-containing chow or control chow for three weeks. Mice were fasted for 18 hours then refed 3 hours prior to sacrifice. Mice were lightly anesthetized with isoflurane. Blood was collected into a microcentrifuge tube containing DPP-IV inhibitor (Millipore Cat #: DPP4, DPP4-010) immediately after sacrifice by decapitation. Serum was collected according to manufacturer’s protocol. Total and active GLP-1 concentrations were measured according to manufacturer’s protocol (Total GLP-1; Millipore GLP-1 Total ELISA Kit Cat #: EZGLP1T-36K, Active GLP-1; Millipore High Sensitivity GLP-1 Active Chemiluminescent ELISA Kit Cat #: EZGLPHS-35K). Data were fit to standards using a five-parameter logistic function.

### Statistical Analysis

Statistical analyses assessing potential sex-specific effects were performed using two-way analysis of variance (ANOVA). Three-way ANOVA was used to assess the contributions of sex, intestinal region, and canagliflozin treatment. Each region of the small intestine was treated as a repeated measurement. Three-way ANOVA of adenoma size analyzed data points from individual polyps grouped within the random effect of mouse ID. Differences within each sex and region were assessed by two tailed Student’s t-test, and P-values were adjusted for multiple comparisons using the Holm-Sidak method. Three-way ANOVA was used to assess the contributions of sex, canagliflozin, and neoplastic vs. normal tissue to protein expression in immunohistochemical analyses. Two-way ANOVA was used to assess the impact of sex and canagliflozin on total and active GLP-1 levels. Three-way ANOVA was used to assess the relative contributions of sex, canagliflozin, and adenoma vs. normal tissue to protein expression in the ileum. Normal distribution of data was assumed. P-values < 0.05 were considered significant. Data were plotted with mean and standard error of the mean (SEM).

## Acknowledgements

Supported by NIA grants U01AG022303 and R01GM101171. Drs. E. Fearon and S. Pletcher are acknowledged for helpful discussions.

## Author contributions

DBL, RAM, and JK conceptualized the study; JK, MES, and JB conducted experiments; JK, MES, RAM, and JKG performed data analysis; JK, DBL, and RAM wrote the manuscript.

## Supplementary Figures

**Supplementary Figure 1.**
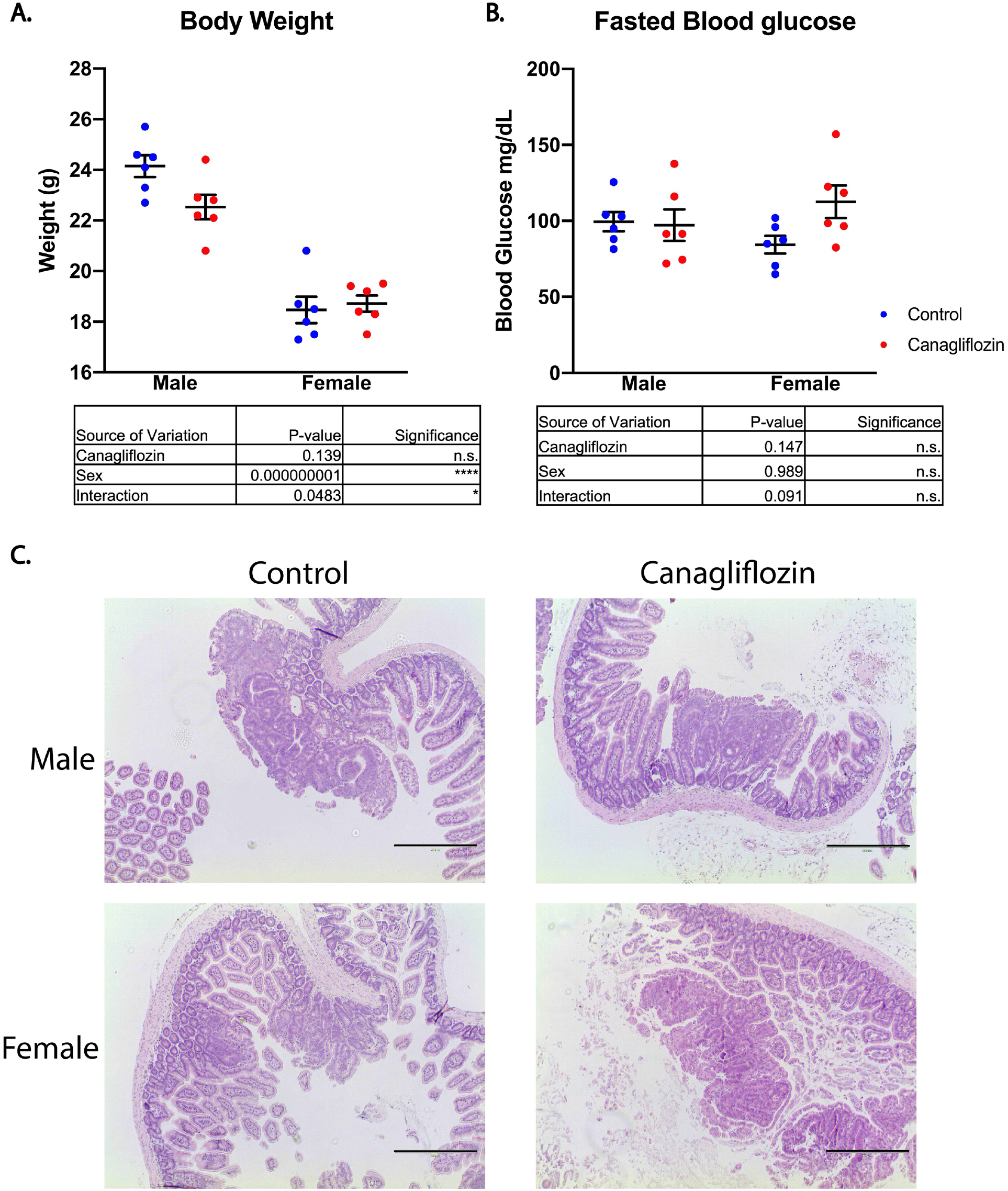
**A)** Body weight measurements of APC^Min/+^ mice were performed immediately after sacrifice. Each data point represents one mouse. Data were analyzed by 2-way ANOVA; * p < 0.05, ** p < 0.01, *** p < 0.001, **** p < 0.0001. **B)** Blood glucose was measured from a tail tip incision of APC^Min/+^ mice using a glucometer. Each point represents the mean of two technical replicates from one mouse. Data were analyzed by 2-way ANOVA. **C)** Representative images of H&E staining of ileal polyps. Scalebar = 400 μm.

**Supplementary Figure 2.**
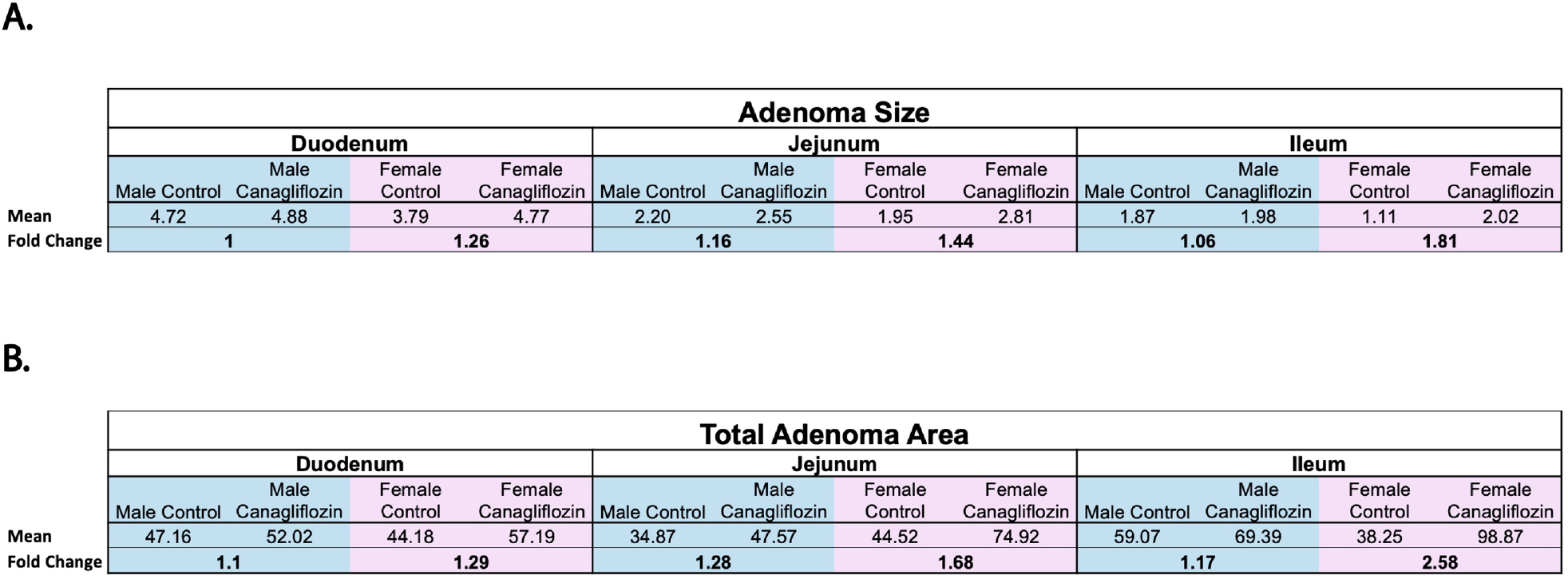
Mean values of **A)** adenoma size and **B)** adenoma area from each region of the small intestine. Fold change between control and canagliflozin groups are displayed separately for males and females.

**Supplementary Figure 3.**
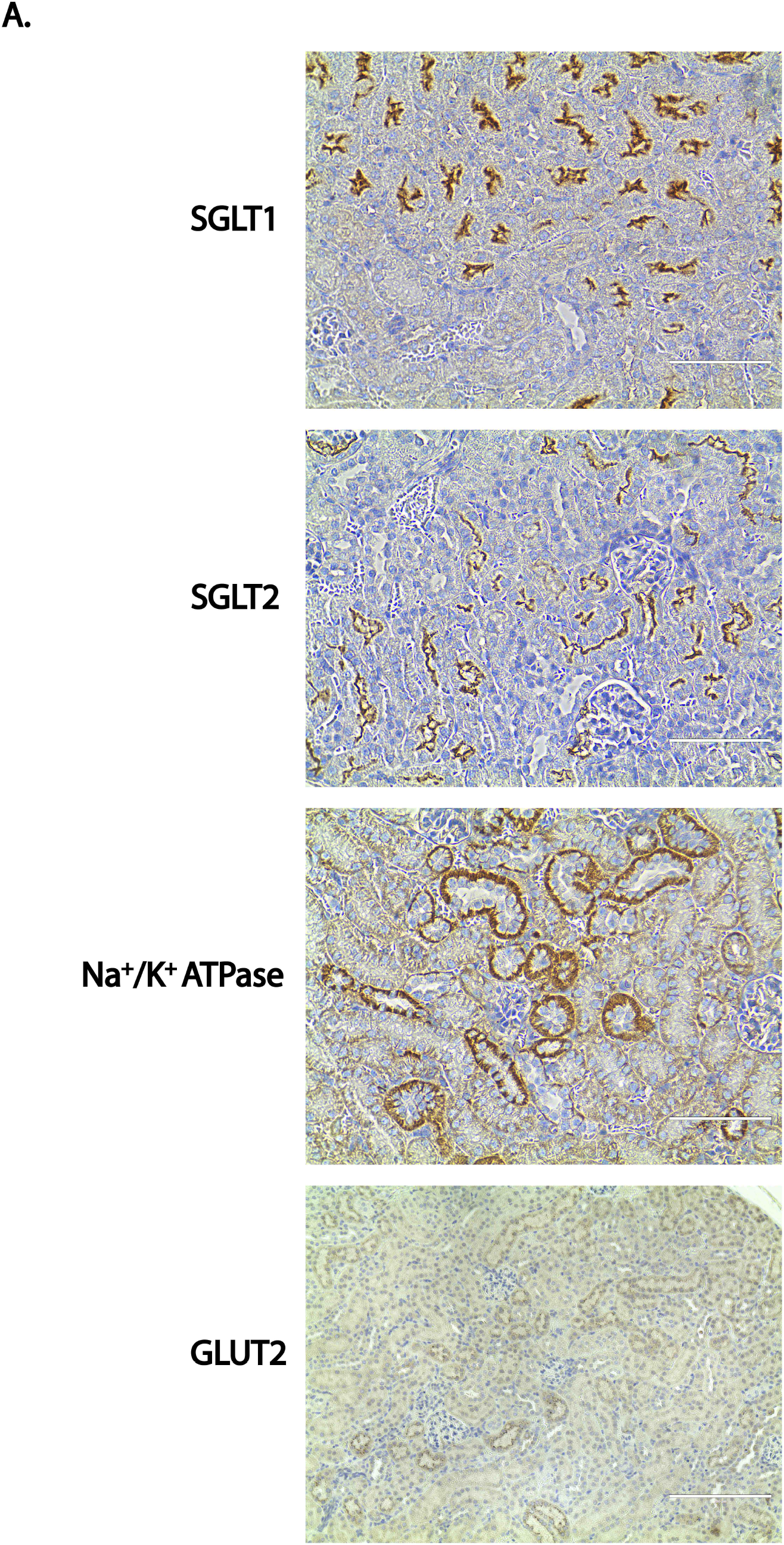
Positive controls for immunohistochemical staining in mouse kidney. SGLT1, scalebar = 100 μm; SGLT2, scalebar = 100 μm; Na^+^/K^+^ ATPase; scalebar = 100 μm; GLUT2, scalebar = 200 μm.

